# A tale of two tails - efficient profiling of protein degraders by specific functional and target engagement readouts

**DOI:** 10.1101/2020.09.22.307926

**Authors:** Alexey L. Chernobrovkin, Cindy Cázares-Körner, Tomas Friman, Isabel Martin Caballero, Daniele Amadio, Daniel Martinez Molina

## Abstract

Targeted protein degradation represents an area of great interest, potentially offering improvements with respect to dosing, side effects, drug resistance and reaching ‘undruggable’ proteins compared to traditional small molecule therapeutics. A major challenge in the design and characterization of degraders acting as molecular glues is that binding of the molecule to the protein of interest (PoI) is not needed for efficient and selective protein degradation, instead one needs to understand the interaction with the responsible ligase. Similarly, for proteasome targeting chimeras (PROTACs) understanding the binding characteristics of the PoI alone is not sufficient. Therefore, simultaneously assessing the binding to both PoI and the E3 ligase as well as the resulting degradation profile is of great value. The Cellular Thermal Shift Assay (CETSA) is an unbiased cell-based method, designed to investigate the interaction of compounds with their cellular protein targets by measuring compound-induced changes in protein thermal stability. In combination with mass spectrometry (MS) CETSA can simultaneously evaluate compound induced changes in the stability of thousands of proteins. We have used CETSA MS to profile a number of protein degraders, including molecular glues (e.g. IMiDs) and PROTACs to understand mode of action and to deconvolute off-target effects in intact cells. Within the same experiment we were able to monitor both target engagement by observing changes in protein thermal stability as well as efficacy by simultaneous assessment of protein abundances. This allowed us to correlate target engagement (i.e. binding to the PoI and ligases) and functional readout (i.e. degrader induced protein degradation).

## Introduction

The mainstream in drug discovery is developing chemical modulators of various protein targets. Chemical modulators, typically small molecules that bind to an enzyme or receptor active site, function by locking their target protein in a state that augments or prevents it from performing its intended function. However, around 80% of the human proteome is estimated to lack favorable features for such drug development schemes, i.e. considered undruggable ^1^ and therefore strategies for modulating the function of the bulk of the proteome is warranted. A possible approach to such undruggable proteins has been to regulate target protein abundance or availability, this has been achieved in a preclinical setting by means of gene editing (CRISPR/Cas9) ^2^, blocking transcription ^3,4^, or selectively destroying target proteins. The latter approach gained substantial attention recently due to the increased understanding of the cellular protein disposal machinery via the ubiquitination – proteasome pathway ^1,5–7^. So far, only a handful of such modalities have reached clinical trials, albeit without any reports on the clinical outcome.

The ubiquitin proteasome system consists of a cascade of enzymes that ultimately results in the conjugation of the 76-amino acid protein ubiquitin onto lysine residues of a target protein. Because ubiquitin contains several lysine residues itself, a chain of polyubiquitin can be assembled: chains coupled to different ubiquitin lysine residues act as recognition motifs for various processes. Most apropos, certain linkages send the protein to the 26S proteasome, a large protease complex that recognizes, unfolds, and degrades ubiquitinated proteins. The molecular determinants of which proteins are targeted for ubiquitination are defined by a class of enzymes known as E3 ubiquitin ligases. Ubiquitin ligases are comprised of several protein subunits with enzymatic functions that result in ubiquitination of a target protein, but the ligase also needs non-enzymatic subunits for target protein recognition in order to ensure a regulated breakdown of proteins ^6^. The specificity of the substrate recognition subunit can be modulated with small molecules, specifically with the phthalimide class of compounds known as immunomodulatory drugs (IMiDs). First described in the 1950s and 1960s as a sedative, the IMiD thalidomide was later discovered to be a potent teratogen ^8^ causing severe phocomelic side effects to newborns. However, since then, thalidomide and its close analogs have been repurposed as potent anticancer agents ^9,10^. Given the excitement regarding thalidomide’s anticancer activity, particularly for multiple myeloma, medicinal chemistry efforts were undertaken to develop related compounds preserving these beneficial activities ^11^. In 2010, a major step towards understanding IMiD action was made when cereblon (CRBN) was identified as a major target driving the thalidomide teratogenicity ^12^. Using a chemoproteomic probe of immobilized thalidomide, the authors identified CRBN as a substrate adapter for the CRL4a ubiquitin ligase complex, whose auto-ubiquitination is inhibited by thalidomide. Importantly, mutations in CRBN that block thalidomide binding also abolish thalidomide teratogenicity in chicken and zebrafish ^12^. Another major breakthrough in the understanding of IMiDs was the identification of the transcription factors Ikaros and Aiolos as proteins that were ubiquitylated upon IMiD treatment ^13–16^. Whereas IMiDs decrease CRBN auto-ubiquitination, they also increase Ikaros ubiquitination. Similarly, cells expressing an IMiD-resistant Ikaros mutant (Q147H) are resistant to IMiD-induced cytotoxicity ^13,15^. More recently, casein kinase 1α (CK1α) was also identified as a protein selectively degraded by the IMiD(/thalidomide analogue) lenalidomide ^17,18^. Finally, a number of proteome-wide studies extended the list of IMiDs-induced neosubstrates of CRBN ^16,19,20^.

Although IMiDs have found clinical success, the applicability of the system is currently limited. As mentioned before, rationally designing a thalidomide analog to target specific proteins for degradation would be difficult given the small structural determinant on the potential substrate, making it challenging to predict and exploit. A strategy to go around this problem was presented by Sakamoto and colleagues that utilized bi-functional molecules, in this case comprised of a small molecule binding the target protein MetAP2 and a peptide that bound the SCF-β-TRCP E3 ubiquitin ligase. This bifunctional molecule was successful in bringing MetAP2 in close proximity to the E3 ubiquitin ligase, which resulted in increased MetAP2 ubiquitination and subsequent degradation ^7^. These types of molecules were later denoted PROTACs (Proteolysis Targeting Chimeras). Also IMiD molecules have been adopted as E3 ubiquitin ligase binding domains and selective ubiquitination and degradation of PoIs was achieved when IMiDs were coupled via a specific linker to a PoI-binding moiety ^1,5,21^. An example of such a molecule is BSJ-03-204 that comprises Pomalidomide as the CRBN-binding domain and Palbociclib as the PoI-binding domain^22^. The main PoIs, in this case CDK4 and CDK6, were ubiquitinated and degraded upon administration of BSJ-03-204 to live cells^22^. Even if the degradation profile of a PROTAC is easy to observe it reports very little about its target engagement profile. The bifunctional nature of a PROTAC molecule could afford a broader target engagement profile than the sum of its components. The opposite is also possible, i.e. that the different domains could hamper each other’s interactions, and as a result decrease the strength and number of interactions. It would therefore be interesting to not only elucidate the quantitative changes in the proteome following PROTAC administration to live cells, but also map the molecule’s high affinity binding partners in both intact cells and lysates.

A crucial part of any drug development program is to ensure proper and specific target engagement and several strategies can be employed to investigate this matter ^23^. The Cellular Thermal Shift Assay (CETSA) is among those methods that can be used in order to investigate target engagement. CETSA is based on the same principle as classical thermal shift assays with the addition that it can be employed in more complex biological systems such as living cells, tissues, whole blood and lysates^24^. Taking advantage of the quantitative mass-spectrometry (MS) based proteomics, CETSA MS allows for the unbiased profiling of a molecule’s interactions and downstream effects on the global proteomic scale (thermal proteome profiling, or TPP), rendering it a powerful tool for target deconvolution and assessment of target specificity of a molecule ^25,26^. We set out to employ CETSA MS and quantitative proteomics to study both the target specificity of several IMiDs in intact cells and cell lysates in order to deconvolute target specificity of IMiDs and to investigate the degradation specificity of IMiDs by quantitative proteomics in intact cells of different origin: transformed human cells (K562); iPSC and iPS derived embryoid bodies (EBs). Lastly, we also compared the observed effects of IMiD treatment with the target engagement and degradation specificity of an IMiD-based PROTAC (BSJ-03-204).

## Materials and Methods

### Cells

K562 cells were obtained from ATCC and were cultured following ATCC protocols. Cell cultures were kept sub-confluent and the cells were given fresh medium one day prior to harvesting. For the experiment, the cells were washed and detached with 1x TrypLE (Gibco, Life Technologies). The detached cells were diluted into Hank’s Balanced Salt solution (HBSS), pelleted by centrifugation (300 g, 3 minutes), washed with HBSS, pelleted again and re-suspended in the experimental buffer (20 mM HEPES, 138 mM NaCl, 5 mM KCl, 2 mM CaCl_2_, 1 mM MgCl_2_, pH 7.4) constituting the 2x cell suspension. For the lysate experiments, K562 cells were re-suspended in the experimental buffer and cells were lysed by three rounds of freeze-thawing. The lysate was clarified by centrifugation at 30 000 x g for 20 minutes and used immediately in the CETSA experiment constituting the 2x lysate. Reagents were purchased from Sigma unless otherwise noted. Cellartis® human iPSC line 22 (ChiPSC22) was obtained from Takara, expanded and cultured using Cellartis DEF-CS Culture System following Cellartis protocol.

Embryoid bodies (EBs) were obtained by seeding ChiPSC22 in *in vitro* differentiation medium (IVD) containing Advanced RPMI 1640, B-27 50X, GlutaMAX 100X, PEST 10,000 units/ml of penicillin and 10,000 µg/ml of streptomycin (Gibco, ThermoFisher Scientfic) and 5µM of Y-27632 (Selleckchem) at a density of 5 × 10^4^ cells per well in a 96-well plate (non-treated, V bottom), centrifugation at 400 x g at room temperature for 5 minutes followed by 7 days incubation at 37°C, 5% CO2 and ≥90% humidity. After 7 days, EBs were transferred into coated 6-well plates (DEF-CS COAT, Takara) at a density of 30 EBs per well in IVD medium and harvested after 7 days.

### Compounds

Thalidomide, Lenalidomide and Pomalidomide were purchased from Sigma Aldrich as lyophilized powders and redissolved in DMSO to 30 mM stock solutions. BSJ03-204 was purchased from Tocris (#6938) and redissolved in DMSO to a 10 mM stock solution.

### 2D CETSA MS experiment

The K562 cell suspension was divided into five aliquots and each was mixed with an equal volume of compound solution prepared at 2x final concentration in the experimental buffer (as above). The resulting final concentrations of compounds (Thalidomide, Lenalidomide or Pomalidomide) were 0.03, 0.3, 3, and 30µM; 1% DMSO only was used as control. Incubations were performed for 60 minutes at 37°C with end-over-end rotation. The treated cell suspensions were each divided into ten aliquots that were all subjected to a heat challenge for 3 minutes, each at a different temperature between 44 and 62°C. After heating, protein aggregates were pelleted by centrifugation at 20 000 x g for 20 minutes and supernatants containing soluble fractions were kept for further analysis.

### Compressed CETSA MS experiment in lysed iPSCs and EBs

iPSCs (ChiPSC22) and EBs were lysed by three rounds of freezing in liquid nitrogen and thawing in a water bath (20°C). Cell debris was removed by centrifugation at 30000xg for 20 minutes. Clarified lysate was then divided into eight aliquots and each was mixed with an equal volume of compound solution prepared at 2x final concentration in the experimental buffer (as above). The resulting final concentrations of Pomalidomide were 0.02, 0.2, 2, 6, 20, 60 and 200 µM; 1% DMSO only was used as control. Incubation was performed for 15 minutes at room temperature. After incubation, samples were divided into 24 aliquots and subjected to a heat challenge for 3 minutes, two samples per temperature point, in a temperature range 44-66°C (in increments of 2°C). After the heating step, samples corresponding to the same compound concentration were pooled together and 16 resulting samples (8 concentration points in two replicates) were centrifuged for 20 minutes at 30 000 x g to pellet aggregated proteins. Supernatants containing soluble fractions were kept for further analysis.

### Time-course Compressed CETSA MS experiment in intact iPSCs and EBs

iPSCs and EBs were incubated with 30 µM of Pomalidomide or vehicle control (1% DMSO) in media (DEF-CS for iPSCs and Advance RPMI and 50X B27 for EBs) for 15, 30 (EBs only), 60, 120 and 480 minutes at 37°C, 5% CO2 and ≥90% humidity. At harvest, the cultures were washed in dPBS [-]CaCl_2_ [-]MgCl_2_, trypsinized with TrplE, washed in dPBS, centrifugated at 200 x g for 3 minutes and resuspended in the experimental buffer (as above). All solutions used for harvest contained 30 µM Pomalidomide or vehicle control, respectively. The cell suspensions were divided into 13 aliquots, of which twelve were subjected to a heat challenge for 3 minutes, each at a different temperature between 44 and 66°C (increment of 2°C), while one aliquot remained unheated. After heating, samples corresponding to the same treatment were pooled together. In all samples, protein aggregates were pelleted by centrifugation at 30 000 x g for 20 minutes and supernatants containing soluble fractions were kept for further analysis.

### LC-MS sample preparation and LC-MS/MS analysis

The total protein concentration of the soluble fractions was measured by DC protein colorimetric assay (BioRad). The same protein amount from each sample was taken and subjected to reduction and denaturation with tris(2-carboxyethyl)phosphine (TCEP) (Bond-breaker, Thermo Scientific) and RapiGest SF (Waters), followed by alkylation with chloroacetamide. Proteins were digested with Lys-C (Wako Chemicals) overnight and trypsin (Trypsin Gold, Promega) for 6 hours.

After complete digestion had been confirmed, samples were labelled with 10-plex Tandem Mass Tag reagents (TMT10, Thermo Scientific) according to the manufacturer’s protocol. Labelling was performed for 3 hours at room temperature and ∼30% ACN in the buffer. Labelling reactions were quenched by addition of hydroxylamine to the final concentration of 0.1%. For the 2D CETSA MS experiment, concentration series from two consecutive temperatures were combined in one TMT10-plex sample. For the eight-concentration point compressed CETSA MS in lysed iPSCs, eight CETSA samples were combined with two non-heated (treated with highest concentration of compound and corresponding DMSO control) within same TMT10-plex sample. For the time-course experiment samples from the same time point were combined together (6 CETSA samples + 4 non-heated samples).

The labelled samples were subsequently acidified and desalted using polymeric reversed phase (Strata X, Phenomenex or Oasis HLB, Waters). LC-MS grade liquids and low-protein binding tubes were used throughout the purification. Samples were dried using a centrifugal evaporator.

For each TMT10-plex set, the dried labelled samples were dissolved in 20 mM ammonium hydroxide (pH 10.8) and subjected to reversed-phase high pH fractionation using an Agilent 1260 Bioinert HPLC system (Agilent Technologies) over a 1 × 150 mm C18 column (XBridge Peptide BEH C18, 300 Å, 3.5 μm particle size, Waters Corporation, Milford, USA). Peptide elution was monitored by UV absorbance at 215 nm and fractions were collected every 30 seconds into polypropylene plates. The 60 fractions covering the peptide elution range were evaporated to dryness, ready for LC-MS/MS analysis.

Thirty individual fractions were analyzed by high resolution Q-Exactive HF or Q-Exactive HF-X mass spectrometers (Thermo Scientific) coupled to high performance nano-LC systems (Evosep One, Evosep, or Ultimate 3000, Thermo Scientific).

MS/MS data was collected using higher energy collisional dissociation (HCD) and full MS data was collected using a resolution of 120 K with an AGC target of 3e6 over the m/z range 375 to 1650. The top 12 most abundant precursors were isolated using a 1.2 Da isolation window and fragmented at normalized collision energy values of 35. The MS/MS spectra (45 K resolution) were allowed a maximal injection time of 120 ms with an AGC target of 1e5 to avoid coalescence. Dynamic exclusion duration was 30 s.

### Protein identification and quantitation

Protein identification was performed by database search against 95 607 canonical and isoform human protein sequences in Uniprot (UP000005640, download date: 2019-02-21) using the Sequest HT algorithm as implemented in the ProteomeDiscoverer 2.2 software package. Data was re-calibrated using the recalibration function in PD2.2. and final search tolerance setting included a mass accuracy of 10 ppm and 50 mDa for precursor and fragment ions, respectively. A maximum of 2 missed cleavage sites was allowed using fully tryptic cleavage enzyme specificity (K\, R\, no P). Oxidation of methionine, deamidation of asparagine and glutamine, and acetylation of protein N-termini were also allowed as variable modification. Carbamidomethylation of Cys, TMT-modification of Lysine and peptide N-termini were set as static modifications.

Peptide-spectrum-matches (PSM) were filtered at 1% FDR level by employing target-decoy search strategy and Perolator rescoring ^27^.

For quantification, a maximum co-isolation of 50 % was allowed. Reporter ion integration was done at 20 ppm tolerance and the integration result was verified by manual inspection to ensure the tolerance setting was applicable. For individual spectra, an average reporter ion signal-to-noise of ≥20 was required. Only unique peptide sequences were used for quantification.

### Data analysis

Protein intensities were normalized ensuring the same total ion current in each quantification channel. Intensity values were then log2-transformed and normalized between treatments and replicates, so the median protein intensity is the same in all treatments and replicates.

The fold-changes of any given protein across the concentration range is quantified by using the vehicle condition as the reference (i.e., a constant value of 1). Fold-changes are also transformed to log2 values, to achieve a normal distribution around 0.

To estimate effect size (amplitude) and p-value (significance) of the protein hits in eight-concentration point compressed CETSA MS experiments, the individual protein concentration-response curve is fitted via *non-linear least squares* algorithm using the following formula:

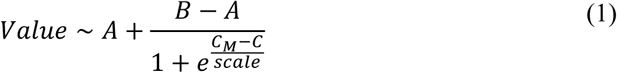

where *Value* — log2-transformed protein intensity, C — log10-transformed compound concentration, A, B, *C*_*M*_ and *scale* — curve parameters for the fit. Using the values from the model fit, the effective concentration corresponding to 50% of maximal signal is estimated: *pEC*_50_ = −*C*_*M*_.

To estimate effect size (amplitude) and p-value (significance) of the protein hits in 2D CETSA MS experiments, protein log-fold changes over Temperature and Concentration domain were fitted via *non-linear least squares* algorithm using the 2D-surface equation formula:

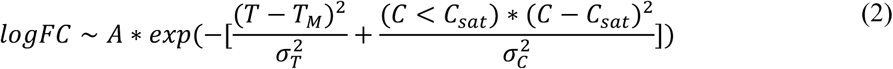

where *logFC*— log_2_-transformed protein fold change relative to the vehicle control at the same temperature, C — log_10_-transformed compound concentration, A, *T*_*M*_, *C*_*sat*_, *α*, and *α*_*c*_— fitting parameters. Using the values from the model fit, the effective concentration corresponding to 50% of maximal signal is estimated: *pEC*_50_ = –*C*_*M*_.

To estimate the significance of time-dependent compound induced changes in either protein abundance or protein thermal stability, following mixed effect a *linear model* was built:

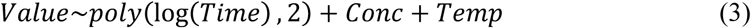

where poly(log(Time),2) – is the second-order polynomic expression using log2-transformed Time value, Conc – two-level factor equal either 1 for Pomalidomide-treated samples or 0 for non-treated samples, and Temp – two-level factor value equal either “CETSA” or “non-CETSA” for heated and non-heated samples respectively.

Significance of the effect for each protein is assessed using ANOVA F-test comparing fit results with the trivial model fit. Benjamini-Hochberg correction is applied to F-test-derived p-values to adjust for multiple comparison.

## Results

Profiling of the three classical IMiDs in intact K562 cells (Figure 1a) was performed using a two-dimensional thermal proteome profiling approach (2D CETSA MS). Here, each protein can be characterized by a heatmap (Figure 1b), where protein fold change (relative to vehicle-control for each temperature) is represented by the color. Alternatively, each protein can be characterized by the observed maximum/minimum log-fold-change (Amplitude) and significance derived from an ANOVA-based F-test to present the results as volcano plots (Figure 1c). Only a handful out of ∼7,000 proteins quantified in the experiments were found to be significantly affected by the IMiDs treatment. Among them we could observe a clear temperature-and concentration-dependent stabilization of CRBN for all three compounds, confirming binding to the endogenous target ^12,28,29^. Notably, for proteins ZFP91, IKZF1 and RNF166 the same concentration dependent reduction of soluble protein amount was observed for all sample temperatures, suggesting compound-induced protein degradation rather than destabilization, which is well in line with previous findings ^14,16,30^. When the same experiment was performed for Pomalidomide in lysed cells (Figure 1c), even fewer proteins showed significant compound-induced thermal stability changes, suggesting that the majority of changes we detected in the intact cells were downstream events, while stabilization of CRBN and glutaredoxin-3 (GLRX3) are caused by direct protein-ligand interactions. Notably, GLRX3 is specifically stabilized only by Pomalidomide, both in intact and lysed cells with a substantial difference in stabilization amplitude (intensity) between intact cells and lysates.

**Figure 1.**
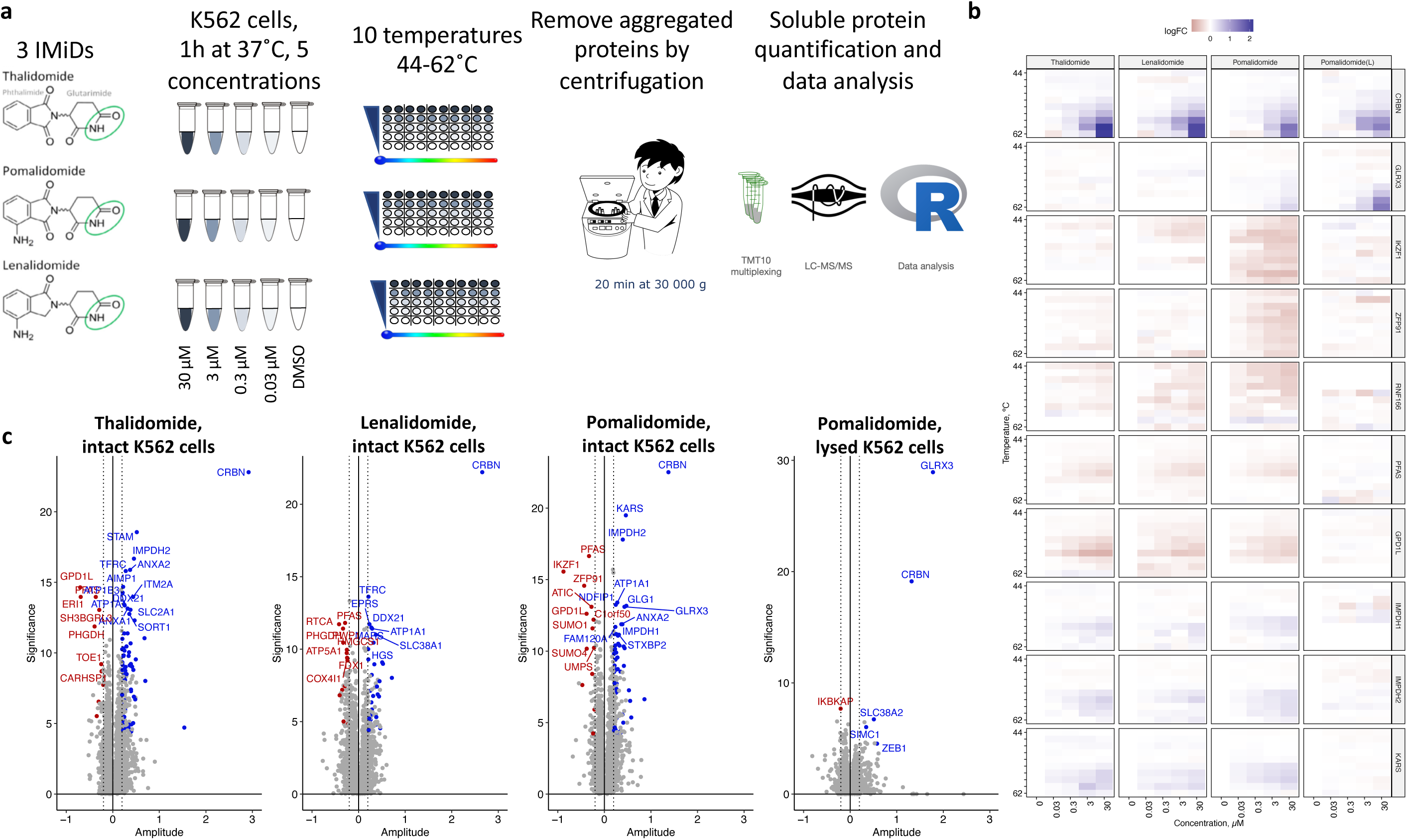
2D CETSA MS profiling of Thalidomide, Pomalidomide and Lenalidomide in intact and lysed K562 cells. (a) an outline of the experiment in intact K562 cells; (b) heatmaps of compound-induced soluble protein amount log-fold changes (relative to untreated control) after heating cell suspensions to different temperatures and removing aggregated proteins via centrifugation; (c) volcano plots of 2D CETSA MS profiling of Thalidomide, Lenalidomide and Pomalidomide in intact K562 cells, and Pomalidomide in lysed K562 cells; x-axis correspond to the CETSA Amplitude (maximum or minimum log-fold change over the heatmap), y-axis corresponds to Significance, which is - log10 transformed p-value from ANOVA F-test (Benjamini-Hochberg adjusted).

The results presented above in the transformed human cell line K562 encouraged us to investigate target engagement and degradation profiles in other sample matrices. In addition to antineoplastic effects, IMiDs are also infamous for their teratogenic effects which are best studied in immature cells. Human induced pluripotent stem cells (iPSC), three-dimensional iPSC aggregates, EBs, in which cells spontaneously differentiate into a heterogenous mix of derivatives of the three germ layers^31^, are the golden standard in *in vitro* teratogenic assessment. iPSCs and EBs were treated with Pomalidomide and target engagement was assessed by CETSA MS and quantitative proteomics was applied to measure protein degradation (Figure 2a).

**Figure 2.**
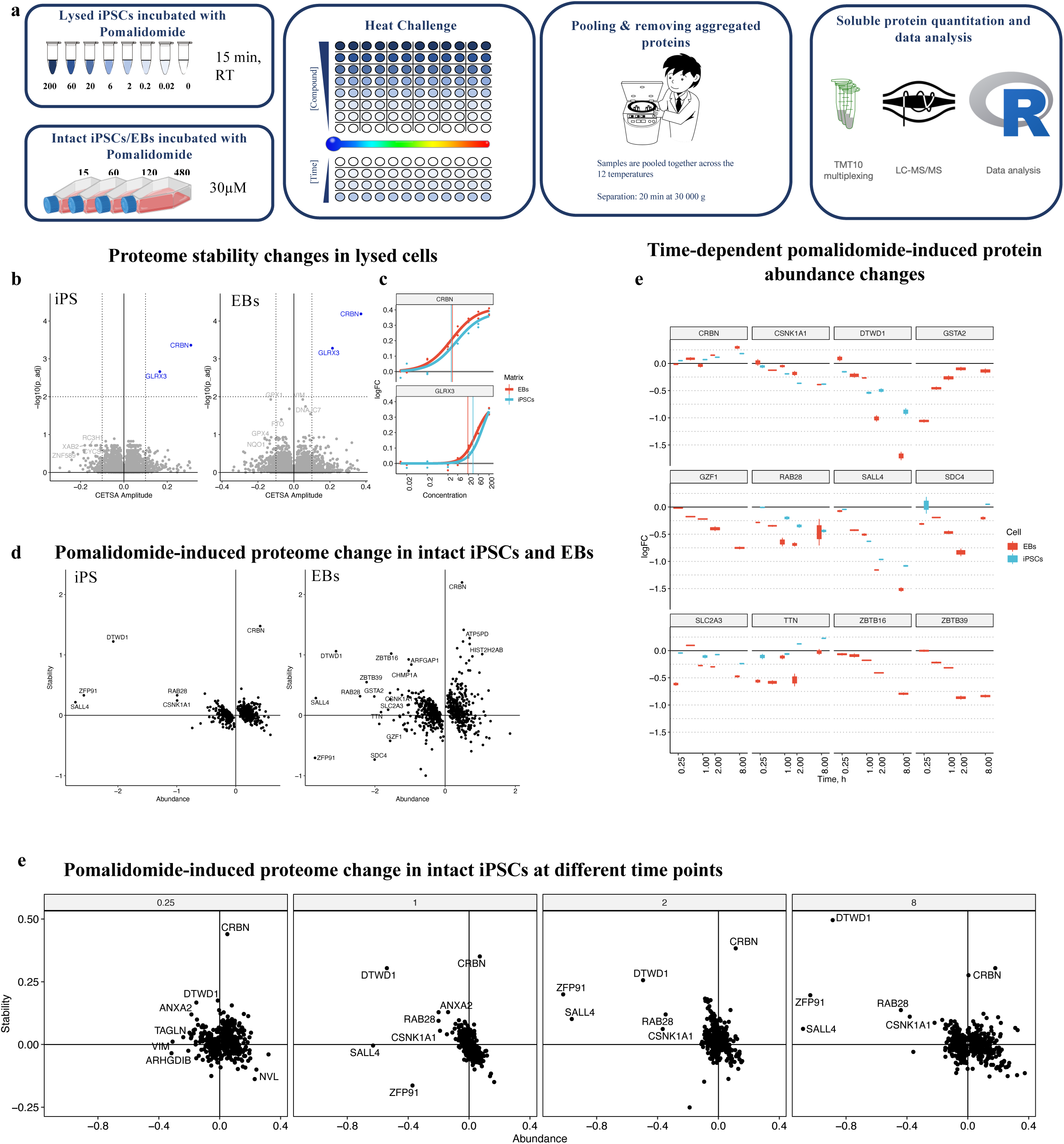
Compressed CETSA MS profiling of Pomalidomide in intact and lysed iPSCs and iPSC-derived EBs. (a) experiment overview – cells/lysate incubation with the compound or vehicle control followed by aliquoting and heat treatment, then pooling over 12 temperatures and removing aggregated proteins by centrifugation; TMT-based LC-MS/MS was used to quantify proteins in soluble fractions; (b) volcano plots of significantly stabilized (positive amplitudes) or destabilized (negative amplitudes) proteins in lysed iPSCs and EBs; (c) concentration-dependent stabilizations of CRBN and GLRX3 in lysed iPSCs and EBs; (d) scatterplots of Pomalidomide-induced stability vs abundance changes in intact iPSCs and EBs; (e) time-dependent compound-induced protein abundance changes in intact iPSCs and EBs; (f) scatter-plot of stability-vs-abundance changes in Pomalidomide-treated iPSCs at different time points.

**Figure 3.**
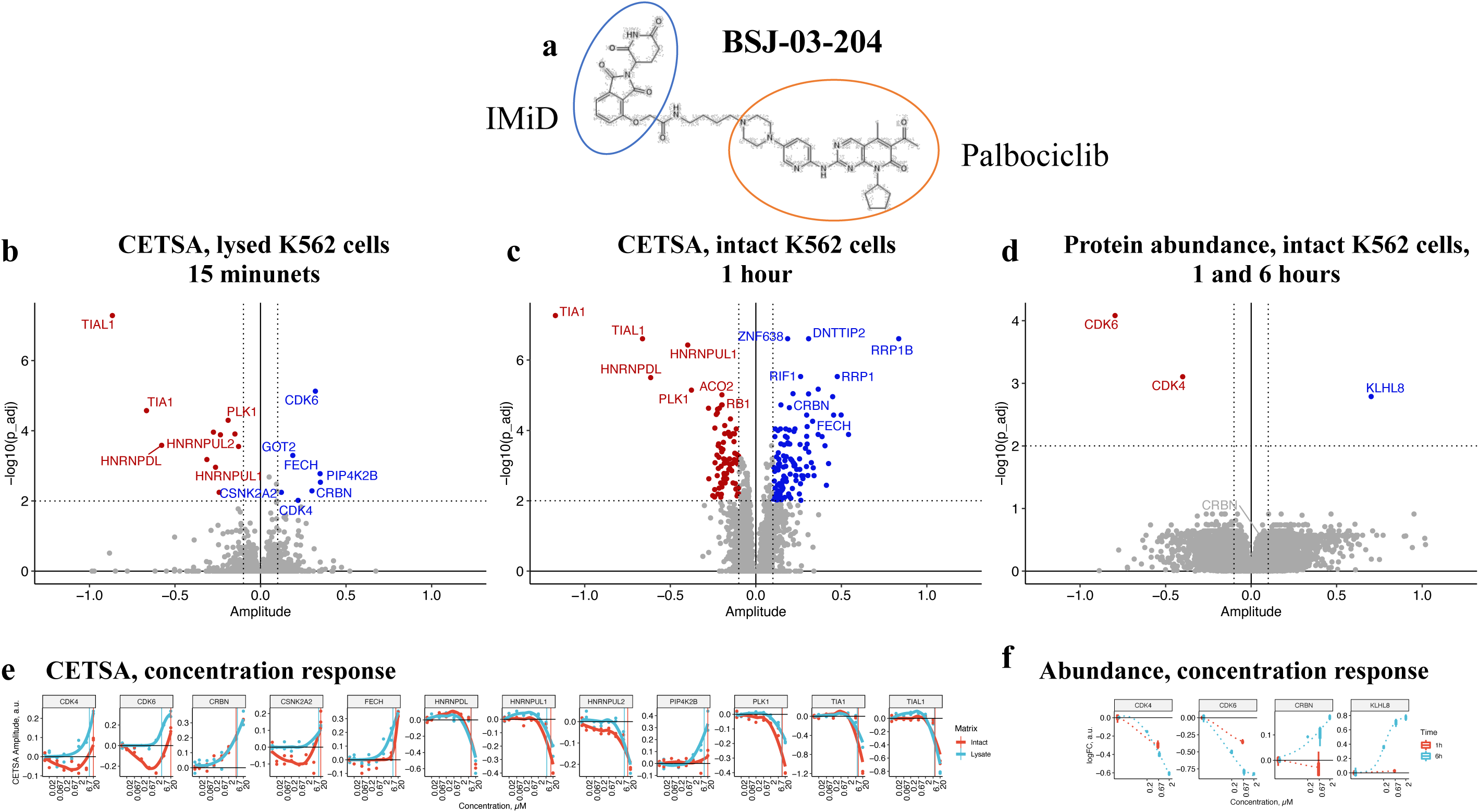
Compressed CETSA MS profiling of PROTAC molecule BSJ-03-204 in intact and lysed K562 cells. (a) chemical structure of BSJ-03-204; (b) volcano-plot of compressed CETSA MS results in lysed K562 cells; (c) volcano-plot of compressed CETSA MS results in intact K562 cells; (d) volcano-plot of time/concentration dependent protein abundance changes in intact K562 cells after 1h and 6h of incubation with different concentrations of BSJ-03-204; (e) compound-induced concentration-dependent protein stability changes in intact and lysed K562 cells; (f) compound-induced protein abundance changes in intact K562 cells after incubation with BSJ-03-204 for 1 and 6 hours.

By applying the recently developed one-pot, or compressed format ^32–34^ we have explored a wider concentration range (lysate experiment), as well as the time-dependence of the binding and degradation profile (Figure 2). Figure 2b shows the results of the eight-concentration Pomalidomide profiling in lysed iPSCs and EBs. In this experiment 7,593 protein groups where quantified. Each protein is represented by its amplitude (relative stability change) and significance (-log10 transformed p-value). CRBN and GLRX3 were the only two proteins that showed a significant concentration dependent change in thermal stability upon treating either lysed iPSCs or EBs with Pomalidomide, supporting the observations in lysed K562 cells (Figure 1c). Notably, stabilization of GLRX3 happens at one-log higher concentrations compared to CRBN (Figure 2c). When treating lysed cells with even higher concentrations of Pomalidomide, we observe changes in stability of more redox-related proteins, including for example stabilization of peroxiredoxins 1,2 and 3, and destabilization of glutathione peroxidases 1 and 4 (Supplementary figure 1).

The previous experiments showed that a single concentration of Pomalidomide of 20-30 µM allows for nearly complete stabilization of cellular targets of Pomalidomide, without causing major off-target binding events. In order to study binding events over time, human iPSC and EBs were incubated with either 30 µM Pomalidomide or vehicle control for 15 min, 30 min (EBs only), 1 h, 2 h, and 8 h. We’ve combined heat-and-pool (CETSA compressed) and non-heated (protein abundance) samples within one TMT-multiplexed sample which allowed us to apply a mixed-effect linear model to the protein measurements to retrieve Pomalidomide-induced Abundance and Stability change contributions into the measured Pomalidomide-induced protein intensity changes (Figure 2c).

In this experiment 7,084 and 8,011 protein groups were quantified in iPSCs and EBs respectively. The higher number of detected protein groups in EBs correlates well with the increased cellular diversity from three germ layers. In line with the 2D CETSA MS results in K562 cells, a clear degradation of the known targets ZFP91, and DTWD1 is observed already after 1 hour of incubation in both iPSCs and EBs cells (Figure 1b, Ikaros protein was not identified in iPSCs and EBs). Additionally, time-dependent degradation of SALL4, RAB28 and CSNK1A1 was observed in two cell systems. Profiling in EBs identified even more Pomalidomide-induced CRBN neosubstrates, including ZBTB16, ZBTB39 and GZF1 (Figure 2c and 2d). Other proteins like GSTA2, SDC4, SLC2A3 or TTN also showed significant alteration of protein abundance, however, the time-course profile of these proteins (Figure 2e) suggest that these are not degradation events. These slightly noisier proteome responses are more pronounced in EB data compared to iPSCs and might reflect the fact that EB samples are heterogenous and prone to have more sample-to-sample variation.

Stabilization of CRBN is observed at all time points studied (Figure 2d). Moreover, time-dependent increase in CRBN abundance can be also seen, likely caused by the inhibition of autoubiquitination. Also, for the majority of CRBN neosubstrates, degradation is accompanied with an increase in protein stability, which is most prominent for the DTWD1. Such protein stabilization has not been detected in lysed cells and might indicate protein stabilization by forming a ternary complex with Pomalidomide and CRBN only in intact cells.

We next explored the effect of the IMiD-based PROTAC BSJ-03-204 on the proteome of K562 cells. In order to evaluate the compound-induced proteome stability changes, intact and lysed K562 cells were incubated with seven concentrations of compound (0.02, 0.067, 0.2, 0.67, 2, 6.7 and 20 µM) for 1 hour and 15 minutes, respectively. The same samples used for intact CETSA MS profiling were also subjected to soluble protein abundance measurements without any heat treatment applied either directly (0, 0.67µM) or after additional 5 hours of incubation (0, 0.2, 0.67 and 2µM) to trace drug-induced protein degradation (Figure 4d, f).

In this experiment heat-treated (CETSA) samples and non-heated samples were measured as different TMT-multiplexed samples. In the heat-treated samples 5,122 and 5,209 protein groups were quantified in lysed and intact K562 cells respectively. Among the proteins that changed stability in lysate were the E3 ubiquitin ligase CRBN and PoIs CDK4 and CDK6, but also other kinases (CSNK2A2, PIP4K2B, PLK1), as well as ferrochelatase (FECH), a common kinase-inhibitor off-target ^35,36^ (Figure 4b). Intriguingly, at first glance no CDKs were found to be affected in the data from intact cells (Figure 4c). This is likely due to the compressed CETSA MS signal consisting of a combination of changes in protein abundance and changes in protein stability. In the intact cell experiment BSJ-03-204 causes both degradation of CDK4 and CDK6 as well as thermal stabilization (as is evident in the lysate experiment). These two effects have opposing results on the protein level in the soluble fraction, i.e. degradation is masked by thermal stabilization and vice-versa. However, when the concentration of BSJ-03-204 is high enough to ligate CDK4/6 and CRBN into separate binary complexes, ubiquitination of the PROTAC substrates is diminished and no degradation occurs (so called “hook effect”, previously described for several PROTACs ^21^) (Figure 4e).

BSJ-03-204 has additional hits in both lysate and intact cells, some of which overlap between sample matrices. TIAL1, TIA1, PLK1, HNRNPUL1 and HNRNPDL were identified as hits in both lysate and intact cell samples, which suggests that these are direct binders to the PROTAC molecule. However, hits that are only occurring in intact cells may constitute pathway effects, induced by the biological effects of CDK4/6 degradation and/or inhibition. One of these proteins with altered thermal stability in response to BSJ-03-204 treatment was RB1 which is a direct substrate of CDK4 and CDK6 ^37,38^. Other proteins affected in the intact cell setting included several proteins involved in transcriptional regulation, RRP1B, RRP1, ZNF638, DNTTIP2, and RIF1. Apart from the stabilization of CRBN, no other similarities were observed between the Pomalidomide and BSJ-03-204 CETSA MS profiles in either intact or lysed K562 cells.

CDK4, CDK6 showed a consistent, time-dependent degradation while CRBN and KLHL8 were rescued from degradation upon the treatment (Figure 4d, f).

## Discussion

In this study we show the target deconvolution capabilities of CETSA MS with regards to delineating the target engagement profiles of the classical IMiDs: Thalidomide, Lenalidomide and Pomalidomide. Moreover, we designed our experiments in such a way that we also could simultaneously monitor both target engagement and pharmacological effect (protein degradation), which resulted in a comprehensive profiling of the investigated IMiDs in relatively short and efficient LC-MS experiments. We took this concept further by applying the same experimental design on the IMiD-based PROTAC BSJ-03-204 enabling us to determine target engagement as well as degradation profiles of BSJ-03-204 in K562 cells.

Target deconvolution by CETSA MS revealed that all IMiDs are relatively selective drugs; CRBN stands out as the high affinity target whose functional modulation is in turn responsible for downstream cellular effects (Figure 1). Other proteins displayed altered thermal stability in intact cells, but only CRBN and GLRX3 did so in lysate. A higher concentration of Pomalidomide was needed to shift GLRX3 compared to CRBN in both K562 and iPSC. In the lysate format most of the biology is turned off, i.e. thermal shifts are more likely to be caused by direct target – ligand interactions and not by signaling events. GLRX3 is an [2Fe-2S] iron-sulphur-complex binding chaperone ^39^, which is interesting when considering another frequent off-target for small molecule drugs: Ferrochelatase (FECH). FECH has also been reported to bind [2Fe-2S] iron-sulphur complexes^40^. We found that BSJ-03-204 stabilized FECH in both intact cells and lysate and FECH has been reported by others as an off-target for many small molecules, including kinase inhibitors ^34–36^. Moreover, even higher Pomalidomide concentrations altered the thermal stability of several other redox-associated proteins. This may be in line with early reports of IMiD’s teratogenic mode of action, in which increased levels of reactive oxygen species were detected upon Thalidomide treatment ^41^.

Three of the proteins that had affected thermal stability in response to IMiD treatment belonged to the inosine synthesis pathway: IMPDH1, IMPDH2, and PFAS. The two formers being the therapeutic targets of the immunosuppressant drug Mycophenolic acid ^42^. Our finding that the thermal stability of IMPDH1 and IMPDH2 were affected downstream of IMiD treatment might account for the immunomodulatory effects of IMiDs. Also, IMPDH expression is increased in neoplasms ^42^, which indicates that the antineoplastic effects of IMiDs could in part stem from modulation of IMPDH function.

We were able to confirm many of the previously identified IMiD neosubstrates in our experiments with iPSC, EBs, and K562 cells. However, not all known neosubstrates were detected in the cell types under investigation. Notably, Ikaros protein (IKZF1) was found to be degraded upon Pomalidomide treatment in immortalized cancer cell line K562 (Figure 1) but was not detected neither in iPSC nor EBs. Aiolos (IKZF3) is a protein related to Ikaros and is also reported to be a neosubstrate ^13,15^, but Aiolos was not identified at all in any of our experiments.

The time dependent degradation profile revealed degradation of the intended targets of BSJ-03-204: CDK4 and CDK6. CRBN on the other hand showed a slight increase that goes in line with the notion that IMiDs inhibit CRBN’s auto-ubiquitination. BSJ-03-204 also had a large impact on the levels of KLHL8 (Kelch-like protein 8) whose levels increased over time. In the nematode (*C. elegans*) a paralog of human KLHL8 (KEL-8) has been reported to function as a substrate adaptor together with Culin 3 (CUL3) containing E3 ligase (BTB-CUL3-RBX1) that controls ubiquitination and degradation of the protein Rapsyn ^43^. Our data suggest that CRBN in turn may regulate ubiquitination and degradation of KLHL8. However, we did not detect the suggested KHLH8 substrate Rapsyn in K562 cells.

When comparing the PROTAC BSJ-03-204 with Pomalidomide alone (Figure 1 and 4) it is striking that there is very little overlap in terms of altered thermal stability and protein abundance, except for the thermal stabilization of CRBN. No degradation of Pomalidomide-induced neosubstrates of CRBN was observed. This in turn indicates that CRBN has been completely hijacked into doing the PROTAC’s bidding, which is degrading CDK4/6.

In conclusion, we have used CETSA MS in combination with quantitative proteomics to study both the induced stability changes of protein targets for different small molecule degraders, as well as the resulting degradation. This has proven to be effective as it allows us to keep track of both target engagement as well as the downstream efficacy. For the IMiDs, our straight-forward approach has allowed us to identify the same primary target as well as downstream preys as have been described collectively during the last 20 years of IMiD research, utilizing a plethora of different techniques. There is currently a large interest in the PROTACs modality and therefore, the need for tools to distinguish between productive binding (i.e those inducing degradation) and silent PROTAC targets, (i.e those that only bind the molecule but don’t form a ternary complex) is much needed.

## Notes

### Competing Interest Statement

DMM is a co-founder and shareholder of Pelago and co-inventor of patents originating from PCT/GB2012/050853. All authors are employees of Pelago Bioscience AB, Sweden. The work was carried with internal funding.

## References

(1) Neklesa, T. K.; Winkler, J. D.; Crews, C. M. Targeted Protein Degradation by PROTACs. Pharmacology and Therapeutics 2017, 174, 138–144. https://doi.org/10.1016/j.pharmthera.2017.02.027.

(2) Zhao, X.; Liu, L.; Lang, J.; Cheng, K.; Wang, Y.; Li, X.; Shi, J.; Wang, Y.; Nie, G. A CRISPR-Cas13a System for Efficient and Specific Therapeutic Targeting of Mutant KRAS for Pancreatic Cancer Treatment. Cancer Letters 2018, 431, 171–181. https://doi.org/10.1016/j.canlet.2018.05.042.

(3) Kole, R.; Krainer, A. R.; Altman, S. RNA Therapeutics: Beyond RNA Interference and Antisense Oligonucleotides. Nature Reviews Drug Discovery 2012, 11 (2), 125–140. https://doi.org/10.1038/nrd3625.

(4) Nijman, S. M. B. Functional Genomics to Uncover Drug Mechanism of Action. Nature Chemical Biology 2015, 11 (12), 942–948. https://doi.org/10.1038/nchembio.1963.

(5) Deshaies, R. J. Prime Time for PROTACs. Nature Chemical Biology 2015, 11 (9), 634–635. https://doi.org/10.1038/nchembio.1887.

(6) Deng, L.; Meng, T.; Chen, L.; Wei, W.; Wang, P. The Role of Ubiquitination in Tumorigenesis and Targeted Drug Discovery. Signal Transduction and Targeted Therapy 2020, 5 (11). https://doi.org/10.1038/s41392-020-0107-0.

(7) Sakamoto, K. M.; Kim, K. B.; Kumagai, A.; Mercurio, F.; Crews, C. M.; Deshaies, R. J. Protacs: Chimeric Molecules That Target Proteins to the Skp1-Cullin-F Box Complex for Ubiquitination and Degradation. Proceedings of the National Academy of Sciences 2001, 98 (15), 8554–8559. https://doi.org/10.1073/pnas.141230798.

(8) Vargesson, N. Thalidomide-Induced Teratogenesis: History and Mechanisms. Birth Defects Research Part C - Embryo Today: Reviews 2015, 105 (2), 140–156. https://doi.org/10.1002/bdrc.21096.

(9) Singhal, S.; Mehta, J.; Desikan, R.; Ayers, D.; Roberson, P.; Eddlemon, P.; Munshi, N.; Anaissie, E.; Wilson, C.; Dhodapkar, M.; Zeldis, J.; Siegel, D.; Crowley, J.; Barlogie, B. Antitumor Activity of Thalidomide in Refractory Multiple Myeloma. New England Journal of Medicine 1999, 341 (21), 1565–1571. https://doi.org/10.1056/NEJM199911183412102.

(10) Vallet, S.; Witzens-Harig, M.; Jaeger, D.; Podar, K. Update on Immunomodulatory Drugs (IMiDs) in Hematologic and Solid Malignancies. Expert Opinion on Pharmacotherapy 2012, 13 (4), 473–494. https://doi.org/10.1517/14656566.2012.656091.

(11) Asatsuma-Okumura, T.; Ito, T.; Handa, H. Molecular Mechanisms of the Teratogenic Effects of Thalidomide. Pharmaceuticals 2020, 13 (5). https://doi.org/10.3390/ph13050095.

(12) Ito, T.; Ando, H.; Suzuki, T.; Ogura, T.; Hotta, K.; Imamura, Y.; Yamaguchi, Y.; Handa, H. Identification of a Primary Target of Thalidomide Teratogenicity. Science 2010, 327, 1345–1350. https://doi.org/10.1126/science.1177319.

(13) Krönke, J.; Udeshi, N. D.; Narla, A.; Grauman, P.; Hurst, S. N.; McConkey, M.; Svinkina, T.; Heckl, D.; Comer, E.; Li, X.; Ciarlo, C.; Hartman, E.; Munshi, N.; Schenone, M.; Schreiber, S. L.; Carr, S. A.; Ebert, B. L. Lenalidomide Causes Selective Degradation of IKZF1 and IKZF3 in Multiple Myeloma Cells. Science 2014, 343 (6168), 301–305. https://doi.org/10.1126/science.1244851.

(14) Lu, G.; Middleton, R. E.; Sun, H.; Naniong, M. V.; Ott, C. J.; Mitsiades, C. S.; Wong, K. K.; Bradner, J. E.; Kaelin, W. G. The Myeloma Drug Lenalidomide Promotes the Cereblon-Dependent Destruction of Ikaros Proteins. Science 2014, 343 (6168), 305–309. https://doi.org/10.1126/science.1244917.

(15) Krönke, J.; Hurst, S. N.; Ebert, B. L. Lenalidomide Induces Degradation of IKZF1 and IKZF3. OncoImmunology 2014, 3 (7). https://doi.org/10.4161/21624011.2014.941742.

(16) Donovan, K. A.; An, J.; Nowak, R. P.; Yuan, J. C.; Fink, E. C.; Berry, B. C.; Ebert, B. L.; Fischer, E. S. Thalidomide Promotes Degradation of SALL4, a Transcription Factor Implicated in Duane Radial Ray Syndrome. eLife 2018. https://doi.org/10.7554/eLife.38430.001.

(17) Petzold, G.; Fischer, E. S.; Thomä, N. H. Structural Basis of Lenalidomide-Induced CK1α Degradation by the CRL4CRBN Ubiquitin Ligase. Nature 2016, 532 (7597), 127–130. https://doi.org/10.1038/nature16979.

(18) Krönke, J.; Fink, E. C.; Hollenbach, P. W.; MacBeth, K. J.; Hurst, S. N.; Udeshi, N. D.; Chamberlain, P. P.; Mani, D. R.; Man, H. W.; Gandhi, A. K.; Svinkina, T.; Schneider, R. K.; McConkey, M.; JärÅs, M.; Griffiths, E.; Wetzler, M.; Bullinger, L.; Cathers, B. E.; Carr, S. A.; Chopra, R.; Ebert, B. L. Lenalidomide Induces Ubiquitination and Degradation of CK1α in Del(5q) MDS. Nature 2015, 523 (7559), 183–188. https://doi.org/10.1038/nature14610.

(19) An, J.; Ponthier, C. M.; Sack, R.; Seebacher, J.; Stadler, M. B.; Donovan, K. A.; Fischer, E. S. PSILAC Mass Spectrometry Reveals ZFP91 as IMiD-Dependent Substrate of the CRL4 CRBN Ubiquitin Ligase. Nature Communications 2017, 8. https://doi.org/10.1038/ncomms15398.

(20) Matyskiela, M. E.; Couto, S.; Zheng, X.; Lu, G.; Hui, J.; Stamp, K.; Drew, C.; Ren, Y.; Wang, M.; Carpenter, A.; Lee, C. W.; Clayton, T.; Fang, W.; Lu, C. C.; Riley, M.; Abdubek, P.; Blease, K.; Hartke, J.; Kumar, G.; Vessey, R.; Rolfe, M.; Hamann, L. G.; Chamberlain, P. P. SALL4 Mediates Teratogenicity as a Thalidomide-Dependent Cereblon Substrate. Nature Chemical Biology 2018, 14 (10), 981–987. https://doi.org/10.1038/s41589-018-0129-x.

(21) An, S.; Fu, L. Small-Molecule PROTACs: An Emerging and Promising Approach for the Development of Targeted Therapy Drugs. EBioMedicine 2018, 36, 553–562. https://doi.org/10.1016/j.ebiom.2018.09.005.

(22) Jiang, B.; Wang, E. S.; Donovan, K. A.; Liang, Y.; Fischer, E. S.; Zhang, T.; Gray, N. S. Development of Dual and Selective Degraders of Cyclin-Dependent Kinases 4 and 6. Angewandte Chemie International Edition 2019, 58 (19), 6321–6326. https://doi.org/10.1002/anie.201901336.

(23) Kubota, K.; Funabashi, M.; Ogura, Y. Target Deconvolution from Phenotype-Based Drug Discovery by Using Chemical Proteomics Approaches. Biochimica et Biophysica Acta - Proteins and Proteomics 2019, 1867 (1), 22–27. https://doi.org/10.1016/j.bbapap.2018.08.002.

(24) Molina, D. M.; Jafari, R.; Ignatushchenko, M.; Seki, T.; Larsson, E. A.; Dan, C.; Sreekumar, L.; Cao, Y.; Nordlund, P. Monitoring Drug Target Engagement in Cells and Tissues Using the Cellular Thermal Shift Assay. Science 2013, 341 (6141), 84–87. https://doi.org/10.1126/science.1233606.

(25) Friman, T. Mass Spectrometry-Based Cellular Thermal Shift Assay (CETSA®) for Target Deconvolution in Phenotypic Drug Discovery. Bioorganic and Medicinal Chemistry 2020, 28 (1). https://doi.org/10.1016/j.bmc.2019.115174.

(26) Savitski, M. M.; Reinhard, F. B. M.; Franken, H.; Werner, T.; Savitski, M. F.; Eberhard, D.; Molina, D. M.; Jafari, R.; Dovega, R. B.; Klaeger, S.; Kuster, B.; Nordlund, P.; Bantscheff, M.; Drewes, G. Tracking Cancer Drugs in Living Cells by Thermal Profiling of the Proteome. Science 2014, 346 (6205), 1255784–1255784. https://doi.org/10.1126/science.1255784.

(27) Käll, L.; Canterbury, J. D.; Weston, J.; Noble, W. S.; MacCoss, M. J. Semi-Supervised Learning for Peptide Identification from Shotgun Proteomics Datasets. Nature Methods 2007, 4 (11), 923–925. https://doi.org/10.1038/nmeth1113.

(28) Lopez-Girona, A.; Mendy, D.; Ito, T.; Miller, K.; Gandhi, A. K.; Kang, J.; Karasawa, S.; Carmel, G.; Jackson, P.; Abbasian, M.; Mahmoudi, A.; Cathers, B.; Rychak, E.; Gaidarova, S.; Chen, R.; Schafer, P. H.; Handa, H.; Daniel, T. O.; Evans, J. F.; Chopra, R. Cereblon Is a Direct Protein Target for Immunomodulatory and Antiproliferative Activities of Lenalidomide and Pomalidomide. Leukemia 2012, 26 (11), 2326–2335. https://doi.org/10.1038/leu.2012.119.

(29) Mori, T.; Ito, T.; Liu, S.; Ando, H.; Sakamoto, S.; Yamaguchi, Y.; Tokunaga, E.; Shibata, N.; Handa, H.; Hakoshima, T. Structural Basis of Thalidomide Enantiomer Binding to Cereblon. Scientific Reports 2018, 8 (1). https://doi.org/10.1038/s41598-018-19202-7.

(30) An, J.; Ponthier, C. M.; Sack, R.; Seebacher, J.; Stadler, M. B.; Donovan, K. A.; Fischer, E. S. PSILAC Mass Spectrometry Reveals ZFP91 as IMiD-Dependent Substrate of the CRL4 CRBN Ubiquitin Ligase. Nature Communications 2017, 8. https://doi.org/10.1038/ncomms15398.

(31) Martin, G. R.; Evans, M. J. Differentiation of Clonal Lines of Teratocarcinoma Cells: Formation of Embryoid Bodies in Vitro. Proceedings of the National Academy of Sciences 1975, 72 (4), 1441–1445. https://doi.org/10.1073/pnas.72.4.1441.

(32) Liu, Y.-K.; Chen, H.-Y.; Chueh, P. J.; Liu, P.-F. A One-Pot Analysis Approach to Simplify Measurements of Protein Stability and Folding Kinetics. Biochimica et Biophysica Acta (BBA) - Proteins and Proteomics 2019, 1867 (3), 184–193. https://doi.org/https://doi.org/10.1016/j.bbapap.2018.12.006.

(33) Gaetani, M.; Sabatier, P.; Saei, A. A.; Beusch, C. M.; Yang, Z.; Lundström, S. L.; Zubarev, R. A. Proteome Integral Solubility Alteration (PISA): A High-Throughput Proteomics Assay for Target Deconvolution. Journal of Proteome Research 2019, 0 (ja), null. https://doi.org/10.1021/acs.jproteome.9b00500.

(34) Chernobrovkin, A.; Lengqvist, J.; Caceres Körner, C.; Amadio, D.; Friman, T.; Martinez Molina, D. In-Depth Characterization of Staurosporine Induced Proteome Thermal Stability Changes. bioRxiv 2020. https://doi.org/10.1101/2020.03.13.990606.

(35) Klaeger, S.; Gohlke, B.; Perrin, J.; Gupta, V.; Heinzlmeir, S.; Helm, D.; Qiao, H.; Bergamini, G.; Handa, H.; Savitski, M. M.; Bantscheff, M.; Médard, G.; Preissner, R.; Kuster, B. Chemical Proteomics Reveals Ferrochelatase as a Common Off-Target of Kinase Inhibitors. ACS Chemical Biology 2016, 11 (5), 1245–1254. https://doi.org/10.1021/acschembio.5b01063.

(36) Savitski, M. M.; Reinhard, F. B. M.; Franken, H.; Werner, T.; Savitski, M. F.; Eberhard, D.; Molina, D. M.; Jafari, R.; Dovega, R. B.; Klaeger, S.; Kuster, B.; Nordlund, P.; Bantscheff, M.; Drewes, G. Tracking Cancer Drugs in Living Cells by Thermal Profiling of the Proteome. Science 2014, 346 (6205). https://doi.org/10.1126/science.1255784.

(37) Nevins, J. R. The Rb/E2F Pathway and Cancer. Human Molecular Genetics 2001, 10 (7), 699–703. https://doi.org/10.1093/hmg/10.7.699.

(38) Knudsen, E. S.; Pruitt, S. C.; Hershberger, P. A.; Witkiewicz, A. K.; Goodrich, D. W. Cell Cycle and Beyond: Exploiting New RB1 Controlled Mechanisms for Cancer Therapy. Trends in Cancer 2019, 5 (5), 308–324. https://doi.org/10.1016/j.trecan.2019.03.005.

(39) Philpott, C. C.; Ryu, M. S.; Frey, A.; Patel, S. Cytosolic Iron Chaperones: Proteins Delivering Iron Cofactors in the Cytosol of Mammalian Cells. Journal of Biological Chemistry 2017, 292 (31), 12764–12771. https://doi.org/10.1074/jbc.R117.791962.

(40) Dailey, H. A.; Finnegan, M. G.; Johnson, M. K. Human Ferrochelatase Is an Iron-Sulfur Protein. Biochemistry 1994, 33 (2), 403–407. https://doi.org/10.1021/bi00168a003.

(41) Parman, T.; Wiley, M. J.; Wells, P. G. Free Radical-Mediated Oxidative DNA Damage in the Mechanism of Thalidomide Teratogenicity. Nature Medicine 1999, 5 (5), 582–585.

(42) Naffouje, R.; Grover, P.; Yu, H.; Sendilnathan, A.; Wolfe, K.; Majd, N.; Smith, E. P.; Takeuchi, K.; Senda, T.; Kofuji, S.; Sasaki, A. T. Anti-Tumor Potential of IMP Dehydrogenase Inhibitors: A Century-Long Story. Cancers 2019, 11 (9). https://doi.org/10.3390/cancers11091346.

(43) Nam, S.; Min, K.; Hwang, H.; Lee, H. O.; Lee, J. H.; Yoon, J.; Lee, H.; Park, S.; Lee, J. Control of Rapsyn Stability by the CUL-3-Containing E3 Ligase Complex. Journal of Biological Chemistry 2009, 284 (12), 8195–8206. https://doi.org/10.1074/jbc.M808230200.

(44) Belair, D. G.; Lu, G.; Waller, L. E.; Gustin, J. A.; Collins, N. D.; Kolaja, K. L. Thalidomide Inhibits Human IPSC Mesendoderm Differentiation by Modulating CRBN-Dependent Degradation of SALL4. Scientific Reports 2020, 10 (1). https://doi.org/10.1038/s41598-020-59542-x.

