## Supplementary figure 1 for "A tale of two tails - efficient profiling of protein degraders by specific functional and target engagement readouts"

### *Simultaneous detection of target engagement and functional readout for in-depth characterization of targeted protein degraders*

Alexey L. Chernobrovkin, Cindy Cázares-Körner, Tomas Friman, Isabel Martin Caballero, Daniele Amadio and Daniel Martinez Molina

#### Supplementary materials

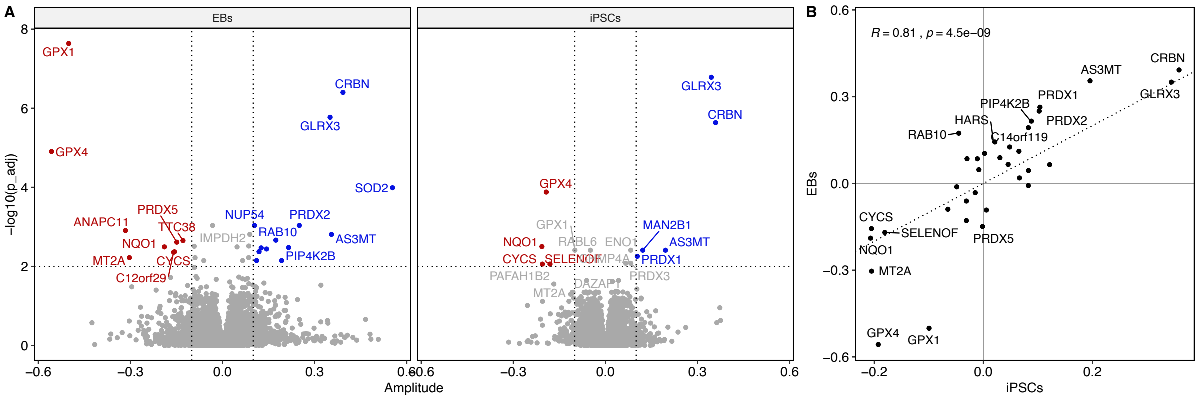


**Supplementary Figure 1. Proteome stability changes in lysed iPSCs and EBs treated with pomalidomide at concentrations up to 200 µM.**
